# Haemolymph removal by the parasite *Varroa destructor* can trigger the proliferation of the Deformed Wing Virus in mite infested bees (*Apis mellifera*), contributing to enhanced pathogen virulence

**DOI:** 10.1101/257667

**Authors:** Desiderato Annoscia, Sam P. Brown, Gennaro Di Prisco, Emanuele De Paoli, Simone Del Fabbro, Virginia Zanni, David A. Galbraith, Emilio Caprio, Christina M. Grozinger, Francesco Pennacchio, Francesco Nazzi

**Author notes:** Corresponding author: Francesco Nazzi.

## Abstract

The association between the Deformed Wing Virus and the parasitic mite *Varroa destructor* has been identified as a major cause of worldwide honey bee colony losses. The mite acts as a vector of the viral pathogen and can trigger its replication in infected bees. However, the mechanistic details underlying this tripartite interaction are still poorly defined, and, in particular, the causes of viral proliferation in mite infested bees.

Here we develop and test a novel hypothesis - grounded in ecological predator-prey theory - that mite feeding destabilizes viral immune control through the removal of both viral ‘prey’ and immune ‘predators’, triggering uncontrolled viral replication. Consistent with this hypothesis, we show that experimental removal of increasing volumes of haemolymph from individual bees results in increasing viral densities. In contrast, we find no support for alternative proposed mechanisms of viral expansion via mite immune-suppression or within-host viral evolution.

Overall, these results provide a new model for the mechanisms driving pathogen-parasite interactions in bees, which ultimately underpin honey bee health decline and colony losses.

## Background

Efficient pollination is vital for crop production (1) and the honey bee is the prevailing managed insect crop pollinator. Honey bees suffer from a range of adverse factors (2); in particular, the Deformed Wing Virus (DWV) is implicated in the substantial colony losses reported in many parts of the world (3). The parasitic mite *Varroa destructor* plays a key role in virus transmission and replication (4, 5); however, there are still alternative and not fully resolved hypotheses about the major mechanisms underpinning the *Varroa-DWV* interaction. The capacity of the *Varroa* mite to transfer DWV was proved by Ball (6) and later confirmed under field conditions (7); these authors also provided preliminary evidence for the replication of the virus within the mite, which was later confirmed (8). However, the mite does not act only as a vector of the virus, thus increasing the pathogen’s prevalence, but can also trigger uncontrolled replication in infected bees, which undermines colony survival (9). Initially, increased replication was attributed to a direct immune suppressive action exerted by the mite (10) but this hypothesis was later questioned (11). It has been suggested that intense replication in infested bees, leading to the development of the characteristic symptom of an overt viral infection, represented by crippled wings at eclosion, is related to the active replication of the virus within the infesting mite (8). However, high DWV copy numbers are frequently detected in bees that are not mite-infested and a clear relationship between viral load within the mites and that in infested bees was not found in all cases (12). Based upon field experiments aiming at assessing the impact of *Varroa* infestation on bees, we showed that the immune challenge represented by the feeding mite amplifies existing viral infections through an escalating bee immunosuppression, perpetuated by the increasing DWV abundance (9). On the other hand, a study describing *Varroa* invasion into a previously mite-free area showed that this was associated with increased DWV prevalence and infection rates, as well as a rapid selection of a single virulent strain, adapted to mite transmission and associated with colony collapse events (13). This facilitation seems to take place also at the individual level when a mite infests a honey bee, where either parasitization or artificial injection favours the replication of a single quasi-clonal DWV strain within the bee (12). In sum, in spite of the large body of evidence about the effect of mite infestation on the dynamics of viral infection in the honey bee and the importance of the *Varroa*-DWV association for honey bee health, there are still alternative and not fully resolved hypotheses on the major underpinning mechanisms. Actually, the available data can also support further and non-contrasting views on how mite feeding can influence the viral titer in bees. In particular, the significant increase in the viral titers of bees infested by three mites versus a single mite (9) and previous observations about the effects of multiple mite infestation on the proportion of symptomatic bees (14) suggests that feeding intensity may play a role. When more *Varroa* mites parasitize the same bee, they make a single wound into the bees’ cuticle to access the haemolymph, and feed from the same opening (15, 16). Thus, the immune response (in particular, melanisation pathways) will be similar for one versus three mites. However, three mites will extract a substantially higher volume of haemolymph from the bee than a single mite, and thus may impact the system by this process. The critical importance of haemolymph removal on DWV dynamics seems to be confirmed by the proliferation of DWV that can be observed after simple wounding with capillary needles and the resulting bleeding from the open wounds (11). Indeed, on a purely theoretical background, it is possible to hypothesize that the concurrent removal of virus particles and circulating antiviral immune effectors by the blood feeding mite can determine a dynamic response similar in principle to that observed when both preys and predators are constantly removed from a predator-prey system (17). Therefore, in order to clarify the causes of viral proliferation in mite infested bees, we carried out controlled lab experiments aiming at testing the existing hypotheses and, in particular, we used both a theoretical analysis and in vivo experiments to test the possibility that mite feeding destabilizes viral immune control through the removal of both viral ‘prey’ and immune ‘predators’, triggering uncontrolled viral replication.

## Methods

### Bees and mites used in this study

The biological material (honey bee worker larvae and *Varroa destructor* adult females) was obtained from an experimental apiary located in Udine (Northeastern Italy). Previous studies indicated that local colonies are hybrids between *Apis mellifera ligustica* Spinola and *Apis mellifera carnica* Pollman (18).

### Artificial infestation of bees with V. destructor mites

The bee larvae and the mites were collected from brood cells capped 0-15 h previously. Larvae obtained as above were transferred into gelatine capsules (Agar Scientific Ltd., 6.5 mm diameter), artificially infested with one mite or left uninfested, and maintained in an incubator (34 °C, 75% R.H., dark) until the adult emergence (19) (Fig. 1A). Newly emerged adult bees and the infesting mites were stored at −80 °C for subsequent molecular analyses. Sixty bees per experimental group were prepared with this method, which, after removing dead bees and mites, resulted in 40 non parasitized honey bees (32 DWV positive) and 32 parasitized honey bees (30 DWV positive) together with 32 infesting mites (27 DWV positive), that were used in the analysis (Fig. 1B). To confirm the results on the distribution of viral infection levels in mite infested bees, another 58 infested bees, prepared as above, were analysed subsequently.

**Fig. 1.**
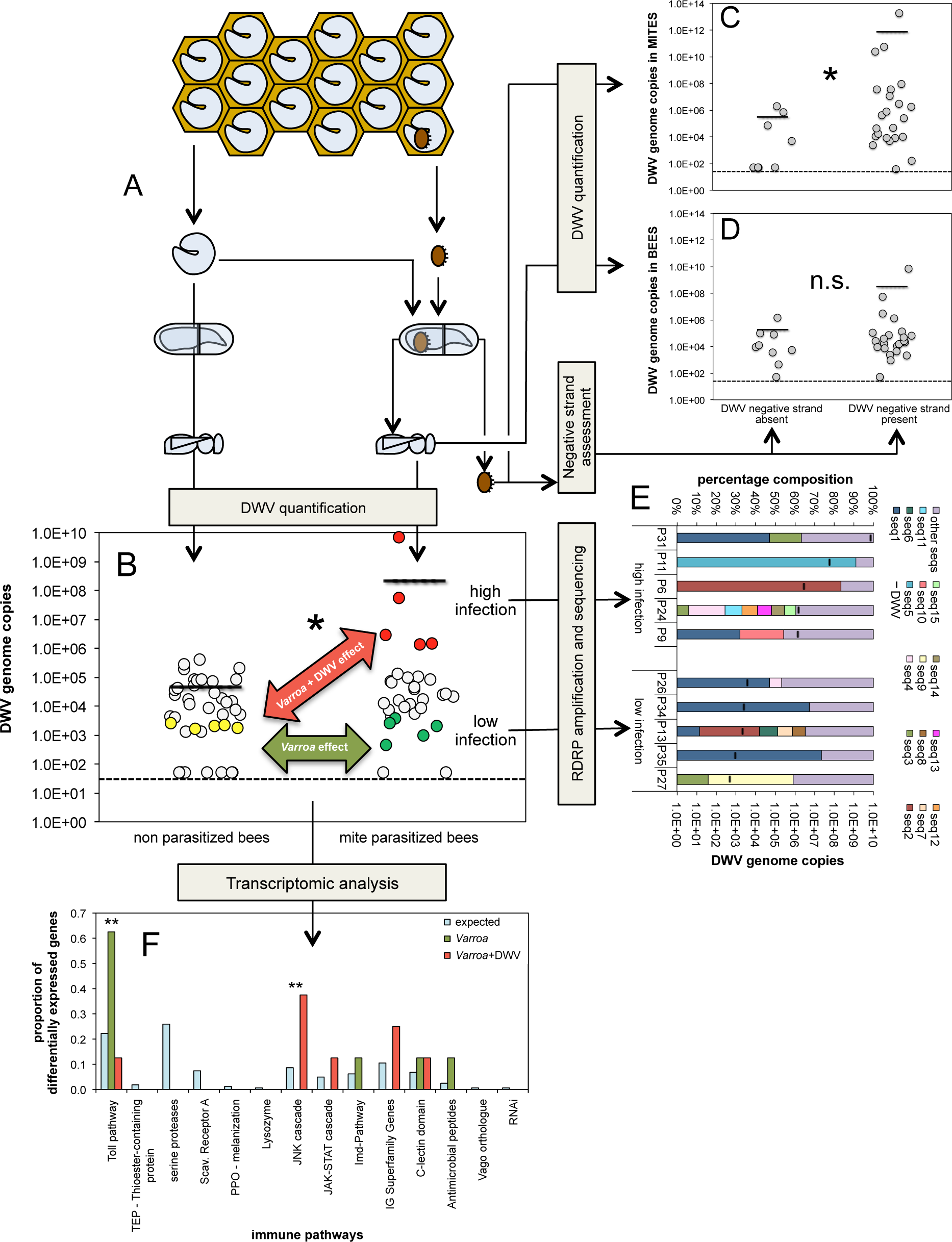
Evaluation of existing hypotheses about the role of *Varroa* mite in increasing virulence of DWV: methods and results. (**A**) Individual bees naturally infected with DWV were artificially infested with one *Varroa* mite or left uninfested. (**B**) Viral load in individual bees infested with one mite or left uninfested as a control. In this and following similar figures, the dashed line represents the lower detection limit for the methodology used; the solid lines represent the average viral load. The samples used for the transcriptomic analysis are marked with different colours: yellow (uninfested-low virus infected bees), green (mite infested-low virus infected bees) and red (mite infested-high virus infected bees). An asterisk marks a significant difference at *P*<0.05. (**C**) DWV genome copies in *Varroa* mites where an active replication was detected (DWV negative strand present) or not (DWV negative strand absent). An asterisk marks a significant difference at *P*<0.05. (**D**) DWV genome copies in bees infested by mites where an active replication was detected (DWV negative strand present) or not (DWV negative strand absent). (**E**) Prevalence of different DWV variants in infected bees with variable virus infection levels. (**F**) Effect of the *Varroa* mite and the combination *Varroa-DWV* on the expression of genes of the canonical immune pathways. The proportion of differentially expressed genes in each pathway, as resulting from the comparison: uninfested-low viral infected bees vs mite infested-low viral infected bees (i.e. *Varroa* effect) and from the comparison: uninfested-low viral infected bees vs mite infested-high viral infected bees (i.e. *Varroa+DWV* effect), is reported as well as the proportion of immune genes belonging to that pathway (i.e. expected). Two asterisks mark significant differences at *P*<0.01 between expected and observed proportions.

In order to test the relative importance of immunity on the increased viral titer normally observed in multiple infested bees, we artificially infested with no, one or three mites five last instar honey bee larvae obtained as described above (Fig. 2A) and compared the expression of immune genes in those bees, as described below.

**Fig. 2.**
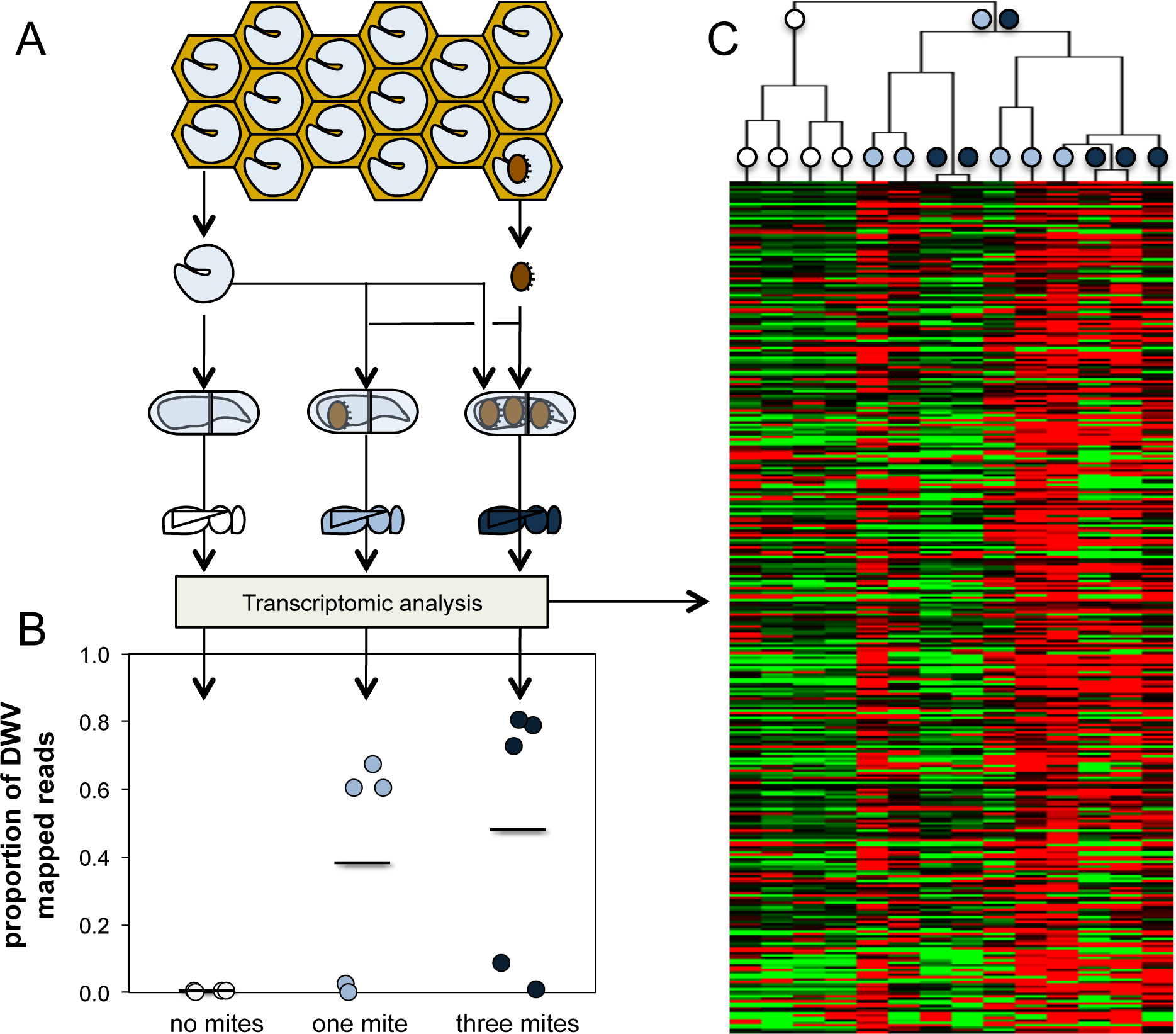
Increased feeding by *Varroa* mite causes increased DWV infection unrelated to immune competence. (**A**) Individual bees naturally infected with DWV were artificially infested with no *Varroa* mites, one mite, or three mites. (**B**) Viral load, as the proportion of reads mapping to DWV genome, in individual bees artificially infested with no mites, one mite, or three mites. (**C**) Clustering of individual bees infested by no mites, one mite, or three mites according to the expression level of immune genes.

### Study of the effects of an increasing haemolymph subtraction on viral replication

This experiment was designed to assess the effect of the removal of an increasing haemolymph volume, in absence of feeding mites, on the dynamics of DWV titer in naturally infected honey bees. Last instar bee larvae were collected from a brood comb as described above and maintained in an incubator (34 °C, 75% R.H., dark) until the white eyes stage, which occurred about 4 days after the collection from brood cells sealed in the preceding 15 hours. Then, 4 experimental groups, made of about 30 pupae each, were established. One group (“control”) was left untreated, whereas all the other bees had the right antenna cut, at the level of the scapum, using fine scissors; pupae bleeding after cutting were discarded. Bees of one group (“wound”) had the wound sealed with a cream containing Sulfathiazole (2%) and Neomycin sulfate (0.5%) to prevent secondary infections. Bees of the remaining two groups (“wound −1 μL” and “wound −2 μL”) had the wound sealed as above, after removing either 1 or 2 μL of haemolypmh, with a microcapillary tube precisely graduated with 1 or 2 μL of ethanol, dispensed through a micropipette; untreated bees acted as further basal control. By subtracting increasing amounts of blood, we tried to assess the effect of pure haemolymph subtraction, while minimizing the impact of wounding and the resulting immune reaction. After treatment bees were kept into a Petri dish, lined with sterile filter paper, and maintained under dark, at 34 °C, 75% R.H., for 4 four days, before assessing the viral titer as described below. To account for the variability across colonies and genotypes, the experiment was repeated four times: on 2 colonies in Udine (Northern Italy) and 2 colonies in Napoli (Southern Italy).

### Quantification of DWV concentration in the honey bee haemolymph

Haemolymph was collected from late instar honey bee larvae by puncturing with an entomological needle the dorsal vessel and collecting the bleeding fluid with a capillary tube. Haemolymph was added to an Eppendorf tube containing the same quantity of anti-coagulant buffer (20), mixed and centrifuged at 1000 g for 10 min at 4 °C. Then, the supernatant containing the plasma was discarded and DWV quantified as described above.

### Molecular analyses

Sequencing of DWV, quantitative DWV analysis, analysis of DWV mutant cloud, DWV Negative strand quantitative analysis and the transcriptomic study of bees were carried out using standard methods described in detail in SI Materials and Methods.

### Statistical analysis of experimental data

The proportion of infected bees (Fig. S2) was compared by means of a Chi Square test; the confidence intervals reported in the figure were calculated with the formula: 1.96√(prop(1-prop)/n). The skewness of the distribution of viral loads across samples was calculated after excluding samples were the virus had not been detected.

Since the normality of the distribution of data about viral infection in bees, as assessed by real time RT-PCR, is not warranted, non-parametric methods were preferred to standard parametric analysis. In particular, the correlation between viral load in mites and bees infested by those mites (Fig. S5) was tested by means of Spearman rank. Comparison between infection levels in bees belonging to different experimental groups was carried out by means of a Mann-Whitney U test (Figure. 1B, 1C, 1D) or Kruskal-Wallis (Figure. 2B, 3). For the statistical analysis of transcriptomic data, see the respective section.

**Fig. 3.**
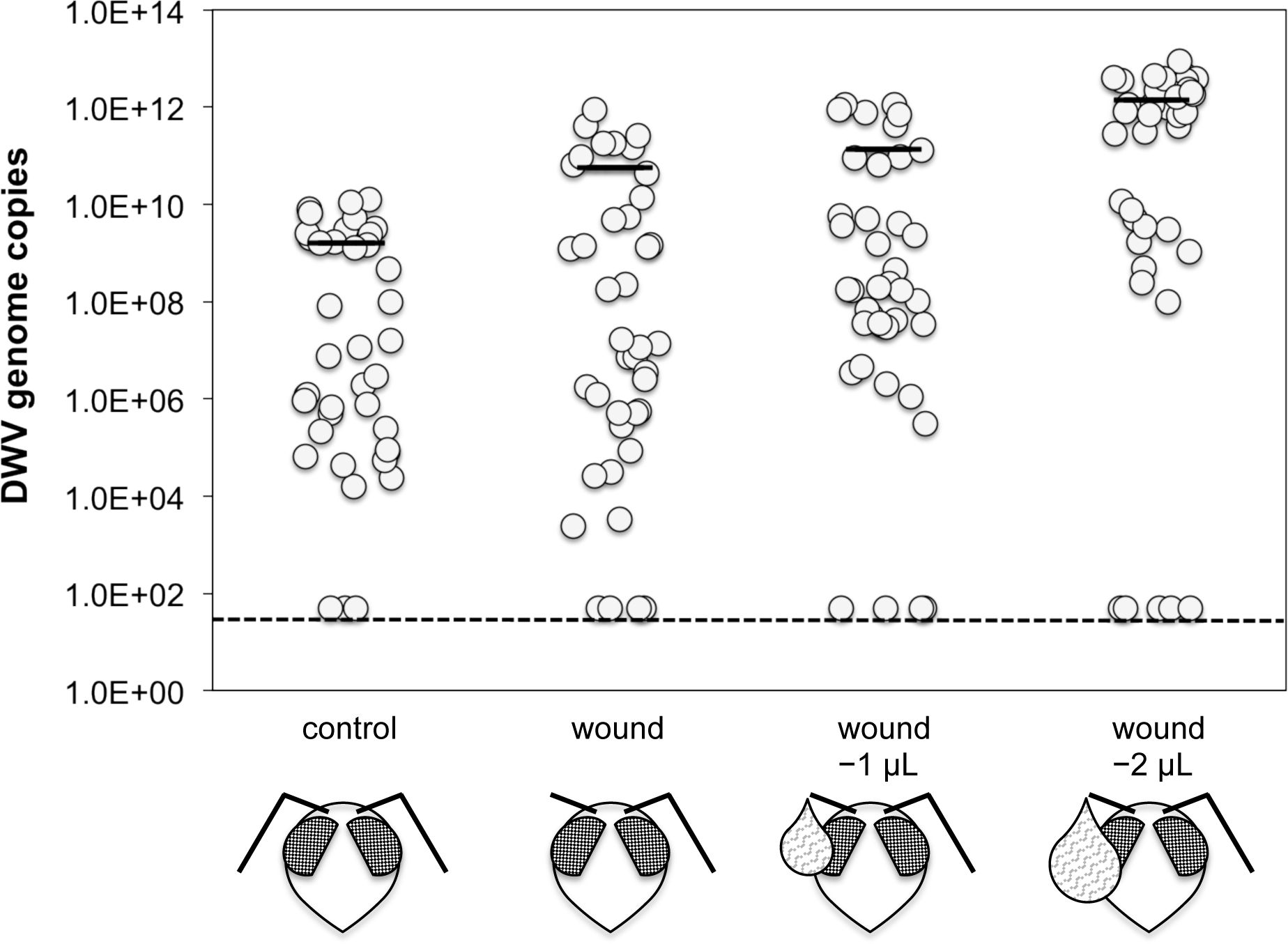
The subtraction of increasing amounts of haemolymph causes increased viral replication in bees. The number of DWV genome copies in bees after the removal of 1 or 2 μL of haemolymph through a wound is reported along the corresponding viral infection in control bees and wounded bees with no haemolymph subtraction.

### Theoretical analysis

In ref. 9 we presented a series of models capturing the coupled within-host dynamics of viral copy number (*V*) and a shared immune currency (*I*). The most parsimonious model analysed further in the current paper included an immuno-suppressive effect of high viral load, as described by the following ordinary differential equations

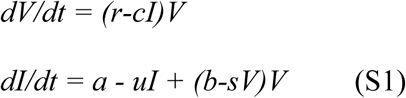

These equations describe the within-host growth of a pathogen population *V* and its controlling immunological counterpart *I*. The maximal rate of pathogen replication is r, which is countervailed by a rate of immunological control *cI*. The dynamics of *I* are shaped by an intrinsic production rate *a*, a rate of decay u, and activation / suppression parameters *b* and *s*. This model implementation ensures that the sign of the impact of virus on immune dynamic (immuno-stimulatory or immuno-suppressive) will depend on viral titre, *V*. Specifically, we assume that at low densities the pathogen is a net activator of immunological activity, whereas at high densities (whenever *V* > *b/s*) the pathogen becomes immuno-suppressive, with *b/s* controlling the threshold point between the two regimes.

To clarify presentation, we again work with a normalized version of equations (S1 in ref. 9) to reduce the parameter dimensions. Specifically, we rescale the units of time to the maximal growth rate of the virus (***t***=*rt*), the units of viral density to the density that halts immune proliferation (***V*** = *Vs/b*) and the units of immune density to the density that halts viral proliferation (***I***=*Ic/r*). Applying these normalizations to equations (S1) lead to the following equations (dropping the bold font for clarity)

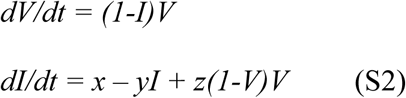

Note that the full system (S1) can be recovered from (S2) by rescaling the units and replacing parameters as follows: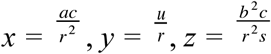. For an equilibrium analysis of equations S2, see ref. 9.

In order to introduce a constant loss of both DWV particles and circulating immune effectors from the system, as caused by the mite feeding upon the bee’s haemolymph, we introduce a rate of *I* and *V* loss *m* (scaled to growth rate r in this dimensionless form) such that the system becomes

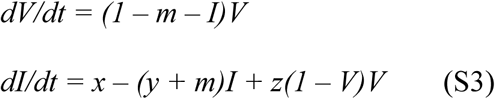

In reference 9 we demonstrate for the *m* = 0 case that there is a lower stable and upper unstable equilibrium in *V*, which converge as *y* increases from zero (see figure 6 in ref. 9). In system (S3) the equilibria become

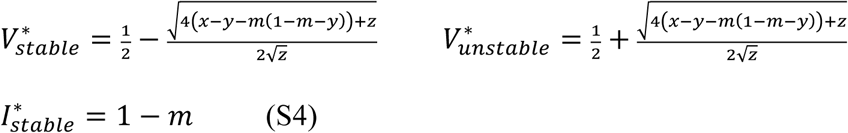

In figure S8 we plot these equilibria as a function of increasing haemolymph removal rate *m*, and observe a decline in immune effectors and an increase in the lower stable equilibrium. To better understand the generality of this effect, we study 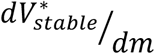, and in the limit of small *m*, we find a tractable result, 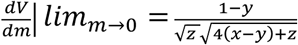, which is positive for reasonable parameter choices – specifically when *y* < 1 and *x* > *y* − *z*/4.

## Results

### Viral infection in parasitized honey bees

To clarify the role of the mite in the dynamics of viral infection in honey bees, we evaluated the presence and abundance of DWV in adult bees that were artificially infested with one mite as mature larvae or were not infested with mites as controls (Fig. 1A); viral presence and titers were evaluated using quantitative real time PCR with sequence-specific DWV primers. Furthermore, a subset of these bees (see Fig. 1B) was subjected to Next Generation Sequencing (NGS) which allowed us to confirm that the bees were infected with DWV and the sequences were >98% identical with a published sequence obtained from a sample collected in the same apiary in 2006 (i.e. NC_004830.2; Fig. S1) and clearly separated from other genotypes of DWV (i.e. NC_006494.1) or recombinants that were associated with higher virulence in other studies (Fig. S1) (12, 21).

We found that 80% of individuals not exposed to mite feeding (n=40) were DWV infected. However, the prevalence of DWV in bees infested by a DWV-infected mite (n=27) was higher at 96% (Fig. S2; Chi Sq.=3.681, d.f=1, P=0.055).

Viral load was higher in bees parasitized by mites compared to control bees (Fig. 1B; average viral load in mite infested bees (n=32)=2.21E+08; average viral load in uninfested bees (n=40)=4.50E+04; Mann-Whitney U=482, n_1_=40, n_2_=32, *P*=0.037). DWV infection levels in non-parasitized bees showed a great variability ranging from 10^˄^3 to 10^˄^6 (Fig. 1B).

However, DWV infection levels showed even greater variability in mite-infested bees; in fact, most mite-infested bees showed infection levels falling within the same interval as that recorded in uninfested bees, but a few specimens largely exceeded the upper limit of this interval, reaching 10^˄^10 viral genome copies per bee (Fig. 1B). Consequently, the distribution of viral loads was very skewed in mite-infested bees (skewness of the distribution of viral loads in mite infested bees (n=30)=5.48, skewness in uninfested bees (n=32)=2.50).

Individual bees sampled later in the field season, when the DWV prevalence and the basal infection rate are higher (9), and artificially infested with one mite, showed a similar skewed distribution of infection levels, with some individuals displaying very high DWV infection levels (skewness of the distribution of viral loads in mite infested bees (n=58)=5.66; Fig. S3). Moreover, re-evaluation of previous data demonstrating the effect of single and multiple mite infestation on viral loads in bees (9) revealed a similar underlying distribution, with a higher median viral infection in mite-infested bees and the distribution of viral loads becoming increasingly sparse (Fig. S4). In sum, the DWV infection data show that the higher viral load observed, on average, in infested bees is due to a change in the distribution of individual viral levels, which is right skewed, due to the presence of a sub-population of highly infected bees as already observed using a different experimental setup (12).

### Viral replication in mites

To study the vector role of *Varroa*, we evaluated the mites infesting the experimental bees above (Fig. 1A) and found that their infection levels were generally higher than those in the bees themselves (median viral load in mites (n=32)= 5.56E+11). A significant correlation was found between the mites’ viral load and viral load of the bees they infested (Fig. S5; n=32, Spearman corr. coeff.=0.531, t=3.433, d.f.=30, *P*=0.002). However, this result cannot be unequivocally interpreted, since the observed correlation could be due either to the fact that a highly infected mite, harbouring an intense viral replication, can inject higher amounts of viral particles, or that a mite infesting a highly infected bee can acquire more virus while feeding.

Active replication of single stranded positive RNA viruses results in the synthesis of the complementary negative strand that is used as a template for the production of viral copies. Therefore, to assess the importance of viral replication within the mite on the level of bee infection, we assessed the presence of DWV negative strands in the mites used for the artificial infestation of bees (Fig. 1A). As expected, the mites containing DWV negative strands had a significantly higher infection level than those where no negative strands were found (Fig. 1C; Mann-Whitney U=42, n_1_=9, n_2_=23, *P*=0.005). However, when we examined whether the viral replication in the parasite was related to the viral load in the host, we found that the infection level of bees infested by mites where an active viral replication was detected was not significantly different from that measured in bees infested by mites which did not apparently harbour an actively replicating virus (Fig. 1D; Mann-Whitney U=80, n_1_=9, n_2_=23, *P*=0.157).

### Composition of the viral mutant cloud

Short replication time and limited correction capability in RNA viruses favour rapid genetic changes, so that, even in a single host, a virus population normally consists of an ensemble of different genetic sequences (i.e. quasi-species). To study the composition of the viral mutant clouds in bees with different levels of viral infection, we amplified and sequenced by NGS the viral region encoding the virus RNA dependent RNA polymerase, by individually sequencing 5 highly infected bees and 5 bees with low infection levels (average DWV genome copies per bee of 1.41E+09 and 1.95E+03, respectively) that were obtained from the previous experiment (see Fig. 1B). From 74 to 559 different variants were reconstructed in each sample, based on a number of viral reads ranging from 40,107 to 160,842. We found no obvious common sequence in low vs high virus infected bees: the most represented sequence was present in six samples from both the low and highly infected groups, at prevalences ranging from 11 to 74% (Fig. 1E). Thus, a link between viral load and molecular diversity was not found (Fig. 1E, Fig. S6).

### Effects of mite infestation and viral infection on the transcriptome of honey bees

To disentangle the effect of the *Varroa* mite parasitization from that of DWV infection on the immune response of bees, we studied the expression of immune genes in bees exposed to a different combination of stress factors (see Fig. 1B; Supplementary Data 1). In particular, to assess the influence of the mite (i.e. *Varroa* effect), we compared the expression level of immune genes in five uninfested bees bearing a low viral infection (average DWV infection=2.04E+03, see Fig. 1B) and five mite infested bees bearing a similar low viral infection level (average DWV infection=1.95E+03, see Fig. 1B). Next, to assess the influence of the combination *Varroa-DWV* (i.e. *Varroa+DWV* effect), we compared five uninfested bees bearing a low viral infection with five mite infested bees bearing a high viral infection level (average DWV infection=1.41E+09). We found that different immune pathways were differentially affected by *Varroa* mite alone and the replicating virus in presence of the mite (Fig. 1F; Supplementary Data 1). Overall, infestation with mites, at low viral infection levels, caused significant changes in expression (i.e. up-regulation) of genes involved in the Toll pathway, while very high DWV infection levels associated with *Varroa* infestation caused significant changes in expression of genes involved in the JNK pathway (Fig. 1F; Supplementary Data 1). Thus, the impact of *Varroa* mite feeding on bee immune response is different from the impact of the high viral titer stimulated by the mite. Furthermore, this experimental setup, allowing the separation of the mite effect from that of the virus, confirmed that immune-suppression by the mites (10) does not play a major role.

### Immune-virus ‘predator-prey’ dynamics within the host

In 2012, we proposed a series of mathematical models describing how within-host viral dynamics are controlled by immunological response, which in turn can be modified by the presence of the virus and other stress conditions, such as mite feeding or pesticide exposure (9, 22). The simplest model consistent with the observation of divergent outcomes (low-cryptic or high-overt infection) required a threshold immune-suppressive effect of DWV. Given this assumption, any factor that depletes the immune system (e.g. increasing mite load) will lead to a gradual increase in a stable DWV set-point until, for sufficiently large depletion, a critical transition to unbound viral replication will follow, leading to overt symptoms and ultimately host death. We hypothesized that, in case of mite infestation, immune depletion may result from the activation of competing immune reactions cross-modulated by shared networks of transcriptional control and, in particular, the melanisation and clotting reactions triggered at the mite’s feeding site, which are under the control of a NF-kB transcription factor that is involved also in antiviral response (9, 23).

Here we mathematically explored the possibility that the concurrent removal of virus particles and circulating antiviral immune effectors by the blood feeding mite can determine a dynamic response similar in principle to that observed when both preys and predators are constantly removed from a predator-prey system (17), by adding to the existing models, which incorporate the depletion of the bee’s immune resources related to the wounding by the parasitic mite (9), a constant rate of loss of both virus and immune effectors, resulting from the feeding of the blood sucking parasite. Indeed, according to this new model, increasing the simultaneous removal of both virus and immune effectors drops immunity and increases/destabilises the viral titre (Fig. S7).

### Effects of the increasing haemolymph subtraction on viral proliferation

In order to verify this hypothesis, we carried out another lab experiment by artificially infesting mature bee larvae with one mite or three mites, using non-infested bees as controls, and by assessing both the viral infection level and immune response at eclosion (Fig. 2A). We observed that the higher DWV titers are associated with the removal of higher amounts of haemolymph, due to heavier mite infestations (Fig. 2B; Supplementary Data 2; Kruskal-Wallis: H=6.41, d.f.=2, *P*=0.041); it is important to note that the lack of a differential immune response in multiple vs single mite infested bees indicates that haemolymph loss alone, rather than an increasing mite-induced immunosuppression, can generate an increasing level of viral infection (Fig. 2C; Fig. S8; Supplementary Data 2; note that no differentially expressed genes were found in the comparison 1 vs 3 mites, whereas 66 and 50 differentially expressed genes were found, respectively, from the comparisons: 0 vs 1 mite and 0 vs 3 mites).

To further corroborate this hypothesis, we assessed the impact of haemolymph subtraction in absence of mite conditioning by comparing viral replication in naturally infected bee pupae from which different amounts of haemolymph were removed with a microcapillary tube from a cut antenna, using wounded or untreated bees as controls. The viral load varied across treatments, with a clear dose-dependent response, positively linking the volume of removed haemolymph to the viral titer measured 4 days after bleeding (Fig. 3; Kruskal-Wallis: H=35.10, d.f.=3, *P*<0.001). In particular, the viral infection in bees to which 2 microliters of haemolymph were removed was about ten times higher than that observed in bees which had only 1 microliter of haemolymph removed from a single wound (Fig. 3; Mann-Whitney U=377, n_1_=n_2_=40, *P*=0.00002), suggesting that blood subtraction alone can play a role, regardless of the immune reaction at the feeding site.

## Discussion

The infection level in uninfested bees is consistent with available data about DWV prevalence in honey bee eggs and larval food (24, 25) and clearly indicates that trans-ovarial and trans-stadial transmission as well as viral acquisition by feeding upon contaminated food during the pre-imaginal life play an important role in the spread of DWV infection within the hive (Fig. S9). The higher proportion of infected bees among those infested by a mite, together with the presence of replicating viruses within the mites, confirms the role of *V. destructor* as a vector of the virus (Fig. S9).

More importantly, our results highlighted the fundamental role of the mite for the increased virulence of DWV in infected honey bees. Collectively, our experimental data allow us to conclude that the capacity of the mite to host the viral pathogen replication (8) (Fig. S10A) appears to be of limited importance for the dynamics of DWV infection in bees. The similar composition and structure of the mutant clouds, observed in low and highly infected bees do not support an important role of viral molecular diversity in the modulation of observed levels of DWV virulence at individual level (Fig. S10B) as proposed earlier (12, 13) but recently questioned (26). Our transcriptomic study further confirms that immune-suppression by the mite (10) (Fig. S10C) does not play an important role. Instead, on the basis of our experimental and theoretical results, we conclude that the stress resulting from mite feeding has the potential of destabilizing the equilibrium between the pathogen and the bee’s immune control (9) (Fig. S10D). Here we confirm our previous hypothesis, based on the depletion of a shared immune resource (9, 23) and show that the intensity of mite feeding can affect the progression of viral infection through a dynamic process triggered by the concurrent removal of the virus and antiviral effectors, which is well described by models proposed for prey-predators interaction (Fig. S10E). The removal of the virus by the feeding mite is a simple consequence of the presence of huge numbers of viral particles in the bee’s haemolymph, as confirmed by the significant correlation between viral infection in bees and the mites which fed upon them, that could not be related to an enhanced within-mite replication. As for the possible presence of antiviral effectors in the bees’ blood and thus the possibility that the mite can subtract significant amounts of them while feeding, it is worth noting that, despite our knowledge about antiviral defence in the honey bee is still incomplete, convincing evidence has been provided regarding the contribute of haemocytes to antiviral defence through phagocytosis in *Drosophila* (27), whereas, according to most transcriptomic analyses, antimicrobial peptides that are dissolved in the bee’s blood certainly play a still uncharacterized role in the immune response to viruses, being constantly up-regulated upon infection (28).

In 1926, the mathematician Vito Volterra, to explain the unexpected fluctuations of certain fish species in the Adriatic Sea, developed his famous model, which clearly showed that the subtraction of both predators and prey, through fishing, could result in the proliferation of the latter (17). Here we suggest, through a modelling approach corroborated by new experimental data, that the pure subtraction of haemolymph - containing both virus and immune factors - from the host, by the feeding mite (Fig. S10E), similarly to the fish industry with regards to prey and predatory fishes, could trigger the proliferation of DWV which can be sustained by the depletion of a shared immune resource (9, 23) and progressively reinforced by the viral induced immune-suppression taking place as soon as the pathogen surpasses a critical threshold (9).

This is a conceptual hypothesis, under which different physiological mechanisms, largely unexplored at molecular level, can fit, and represents the most parsimonious interpretation of the mite role in the enhanced virulence of the virus, providing the logical framework for future experiments aiming to unravel the intimate molecular mechanisms involved.

To our knowledge, this “micro-ecological” perspective of the immune interactions has not been proposed so far for any other blood-feeding parasite and associated pathogens, likely because most species studied from this point of view do not perform a severe bleeding like *Varroa* mites, and, then, the impact on viral dynamics by blood removal is more limited. Indeed, *V. destructor* can consume as much as 0.7 µL of bee haemolymph every 24 h (29), corresponding to the 0.6% of the bee pupa’s haemolymph (30), which, according to our data, can contain up to 10˄3-10˄8 DWV particles per µL according to the infection level.

## Conclusions

We believe that this new information on the interactions within the bee-mite-virus network provides a new vision of the crucial role played by *Varroa* mite in the re-emergence of DWV, an endemic pathogen of honey bees that plays a key role in the current widespread crisis of the beekeeping industry. Moreover, the proposed “micro-ecological” perspective of the immune interactions has broader implications in the research area of animal parasitology.

## Author’s contributions

DA participated in the design of the study, conducted the lab experiments and participated in data analysis; SPB carried out the theoretical analysis and wrote the manuscript; GDP conducted part of the lab experiments and carried out the molecular lab work; EDP, SDF, DAG contributed to data analysis; VZ, EC contributed to lab work; CMG participated in the design of the study and data analysis and wrote the manuscript; FP conceived the study and wrote the manuscript; FN conceived the study, collaborated to lab experiments and data analysis and wrote the manuscript.

## Acknowledgments

We gratefully acknowledge Michelle Flenniken for critically reading an earlier version of this manuscript.

## Supplementary materials and methods

**Sequencing of DWV**. The de novo assembly of DWV was performed using CLC Genomics Workbench 9.0. Briefly, simple contig sequences were first created using the information from the read sequences using an assembly algorithm based on de Bruijn graphs −a compact representation based on short words (k-mers) that is ideal for high coverage, very short read (25-50 bp) data sets- then all reads were mapped using the simple contig sequence as a reference.

A phylogenetic tree was constructed from two sequences obtained in this study (samples P11 and P31, Fig. 1B-1E) and 12 available complete DWV genomes including:

gi|71480055|ref|NC_004830.2| Deformed wing virus, complete genome,

gi|31540603|gb|AY292384.1| Deformed wing virus isolate PA, complete genome,

gi|390190253|gb|JQ413340.1| Deformed wing virus isolate Chilensis A1, complete genome,

gi|592965310|gb|KJ437447.1| Deformed wing virus isolate Varroa-infested-colony-DJE202, complete genome,

gi|40714033|dbj|AB070959.1| Kakugo virus genomic RNA, complete genome,

gi|47177088|ref|NC_005876.1| Kakugo virus, complete genome,

gi|430007848|gb|JX878304.1| Deformed wing virus strain Korea-1, complete genome,

gi|430007850|gb|JX878305.1| Deformed wing virus strain Korea-2, complete genome,

gi|56121875|ref|NC_006494.1| Varroa destructor virus-1, complete genome,

gi|55925812|gb|AY251269.2| Varroa destructor virus 1, complete genome,

gi|301070167|gb|HM067437.1| Deformed wing virus isolate VDV-1-DWV-No-5, complete genome,

gi|301070169|gb|HM067438.1| Deformed wing virus isolate VDV-1-DWV-No-9, complete genome.

Multiple sequence alignment was performed using the ClustalW algorithm and the phylogenetic tree was constructed with “MEGA7: Molecular Evolutionary Genetics Analysis version 7.0 for bigger datasets” (31) using the neighbor-joining method; bootstrap values are based on 1000 replicates.

**Quantitative DWV analysis**. Total RNA was isolated from individual honey bees using TRIzol reagent (Invitrogen), according to the manufacturer’s instructions. The concentration and purity of total RNA were determined by spectrophotometry (Varioskan Flash Spectral Scanning Multimode Reader; Thermo Fisher Scientific).

The quantification of DWV genome copies was performed by SYBR Green qRT-PCR. Titers of DWV were determined by relating the Ct values of unknown samples to an established standard curve. The standard curve was established by plotting the logarithm of seven 10-fold dilutions of a starting solution containing 21.9 ng of plasmid DNA pCR II-TOPO (TOPO-TA cloning) with a DWV insert (from 21.9 ng to 21.9 fg), against the corresponding Ct value, as the average of three repetitions. The PCR efficiency (E=107.5%) was calculated based on the slope and coefficient of correlation (R2) of the standard curve, according to the following formula: E=10(−1/slope)−1 (slope=-3.155, y-intercept=41.84, R2=0.999). Amplifications were performed using the StepOne Real-Time PCR System (Life Technologies) with the following thermal cycling profiles: one cycle at 48 °C for 15’ for reverse transcription, one cycle at 95 °C for 10’; 40 cycles at 95 °C for 15’’, 60 °C for 1’; one cycle at 68 °C for 7’, using the Power SYBR Green RNA-to-Ct 1-Step Kit (Thermo Fisher Scientific). Primer pair (DWV-Forward 5’-GCGCTTAGTGGAGGAAATGAA-3’; DWV-Reverse 5’-GCACCTACGCGATGTAAATCTG-3’) was designed using PrimerExpress 3.0 software (Life Technologies) following the standard procedure. Negative (H_2_O) and positive controls (previously identified positive samples) were included in each qRT-PCR run. According to the manufacturer, the used equipment should allow the detection of the virus provided that at least 50 genome copies are present in the sample.

**Analysis of DWV mutant cloud**. For DWV RNA-dependent RNA polymerase (DWV\RdRp) analysis, specific primer pair were chosen for the amplification of a genome portion of 454 bp (DWV\RdRp-Forward 5’-TAGTGCTGGTTTTCCTTTGTC-3’; DWV\RdRp-Reverse 5’-CCCAGGACCAAAATTCTTAT-3’). The PCR reactions were conducted using the Invitrogen SuperScript III One-Step System (Platinum) following the manufacturer’s procedure. The thermal profile was: 50 °C for 30’; 94 °C for 2’; 40 cycles at 94 °C for 15’’, 60 °C for 30’’, 68 °C for 1’; 68 °C for 5’. Amplified products were run on a 1% agarose gel containing 0.5 μg/ml ethidium bromide and then visualized by UV transillumination. The specificity of the RT-PCR assay was confirmed by sequencing analysis. RT-PCR bands were excised from the agarose gel and purified using the Pure Link Quick Gel Extraction Kit (Invitrogen, Carlsbad, CA). The sequence data of the fragment were analysed using the BLAST server at the National Center for Biotechnology Information, NIH. For the RNAseq analysis, the same primer pairs described above were designed with a specific overhang (DWV\RdRp-Forward 5 ‘-TCGTCGGCAGCGTCAGATGTGTATAAGAGACAG-3’; DWV\RdRp-Reverse 5’-GTCTCGTGGGCTCGGAGATGTGTATAAGAGACAG-3’). All primer pairs were designed by using Primer Express Software (Applied Biosystems).

Raw Illumina paired-end reads in forward and reverse orientation were merged using the PEAR software v0.9.8 (32), adaptor trimmed using cutadapt v1.9 (33) and quality filtered using the FASTX Toolkit’s fastq_quality_filter program (34) requiring minimum quality of 20 in 95% bases or more. For the study of DWV variants we used a software designed for quasi-species reconstruction from next-generation sequencing data (35). Sequenced fragments were aligned against a reference genome and the reference genome was partitioned into a set of sliding windows, then a reconstruction algorithm based on combinations of multinomial distributions was applied. The selected reference genome was the one most closely related to the DWV samples sequenced in this study (i.e. NC_004830.2; Fig. S1). The analysis output is a collection of sequences with prevalences, which was used for a diversity analysis based on the Shannon index.

**DWV Negative strand quantitative analysis**. In order to quantify the DWV negative strand in infesting mites, a strand-specific Biotin/Streptavidin method was used. SYBR-Green real-time quantitative PCR (qPCR) as described above was preceded by a specific retrotranscription incorporated with biotinylated-primers (Biotin-DWV-Forward 5’-TCG ACA ATT TTC GGA CAT CA-3’; Biotin-DWV-Reverse 5’-ATC AGC GCT TAG TGG AGG AA-3’) and magnetic bead purification by using the Dyna beads KilobaseBINDER Kit following the manufacturer’s instructions (Applied Biosystems).

**Transcriptomic study of bees**. Samples were transferred into liquid nitrogen and the total RNA isolated using Tri-reagent (MRC Inc., USA) (Ambion Inc.). RNA sequencing libraries were generated by means of TruSeq Standard mRNA seq kit Illumina, according to the standard protocol indicated by the producer (Illumina, Inc., CA, USA) and starting from 1-2 micrograms of high quality RNA (RNA Integrity Number >7, Agilent Technologies Bioanalyzer, Agilent Technologies, USA). This protocol produced 25-30 million reads per sample that were 36 bases in length.

The sequencing reads were trimmed using ERNE (36) in order to remove low quality reads and the adapters removed with Cutadapt (33). The remaining reads were aligned to the most recent honey bee genome build (Amel 4.5 (37)) with TopHAt2 (38) using default parameters and annotated with the newest official gene set (OGS 3.2). Reads were counted with Htseq count (39) using exons as accepted regions and cumulating the counts at whole gene level.

The obtained data were uploaded into DEseq. (40) to be elaborated by means of VST (Variance Stabilizing Transformation) algorithm and used for the cluster analysis.

Five uninfested and 10 newly emerged infested bees, 5 with a high DWV infection level and 5 with a low infection level, were collected from the cages and processed as above (Fig. 1B). Also five newly emerged bees infested with no, one or three mites (Fig. 2A) were processed as above.

To gain insight into the relative contribution of the *Varroa* mite and DWV in the alteration of the expression of genes belonging to the canonical immune pathways, we studied the transcriptome of bees exposed to a different combination of stress factors (Supplementary Data 1). In particular, to assess the influence of the mite, we compared the expression level of 191 immune genes, listed in ref. 12, in five uninfested bees bearing a low viral infection (average DWV infection=2.04E+03) and five mite infested bees bearing a similar, low viral infection level (average DWV infection=1.95E+03) (Fig. 1B); instead, to assess the influence of the combination Varroa-DWV we compared five uninfested bees bearing a low viral infection with five mite infested bees bearing a high viral infection level (average DWV infection=1.41E+09) (Fig. 1B). Note that 14 and 151 genes classified in ref. 12 as “immune related” and “immune system process” respectively, were not included in this analysis that concentrated on canonical immune pathways.

Due to the uncertainty about the distribution of data, the comparisons were carried out using the Mann-Whitney U test. Since the purpose of this research was not to gather data about single genes but rather to gain an indication about the involvement of different immune pathways, and keeping into account the moderate probability of type I errors, no correction for multiple comparisons was applied but rather differentially expressed genes were considered as such when probability was below 0.01. To verify the possible enrichment of each immune pathway, the proportion of differentially expressed genes belonging to that pathway out of the total number of DEGs in that contrast was compared with the proportion of genes belonging to that pathway out of the total number of genes considered here; Chi square test was used and, again, the level of probability was set to 0.01.

In order to test the relative importance of immunity on the increased viral titer normally observed in multiple infested bees, we artificially infested with none, one or three mites five last instar honey bee larvae obtained as described above and compared the expression of immune genes in those bees, as described above (Supplementary data 2). The proportion of reads mapping onto the DWV genome provided an estimation of the viral infection level in bees whereas a cluster analysis of samples according to the expression of immune genes gave an indication of the effects of an increasing mite infestation on the bee’s immunity.

FPKM were used as gene expression data. The list of honey bee immune genes proposed by Ryabov et al. (12) was used.

To remove possible outlier samples from the analysis, the mean expression value across all samples for each gene was calculated as well as the standard deviation (note that here samples were considered as belonging to a single big group, in order not to introduce any bias in our preliminary filtering). Then expression values higher or lower than the mean expression value ±2 standard deviations were noted. Lastly the proportions of outlier genes in each sample were considered and samples having more than 25% of outlier genes were excluded (this procedure lead to the exclusion of one single sample out of which appeared to have a number of outlier genes bigger by three fold than any other sample).

Then we performed a cluster analysis using “Gene Cluster 3.0” (41) with the following options: centering genes around mean, similarity metrics: correlation (uncentered), clustering method: average linkage. The tree was displayed using “Treeview” (42).

**Figure S1.**
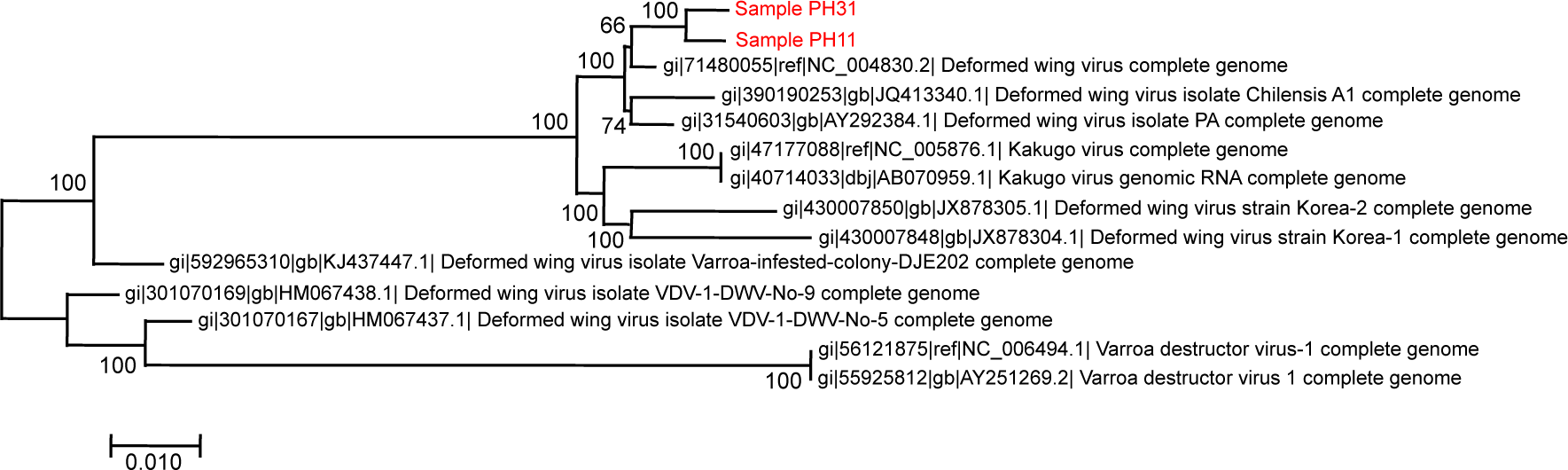
Phylogenetic relationship of the DWV infecting the bees used in this study (samples Pll and P31). The dendrogram is based on the complete genomes (nt sequences) of available sequences of DWV strains and related viruses.

**Figure S2.**
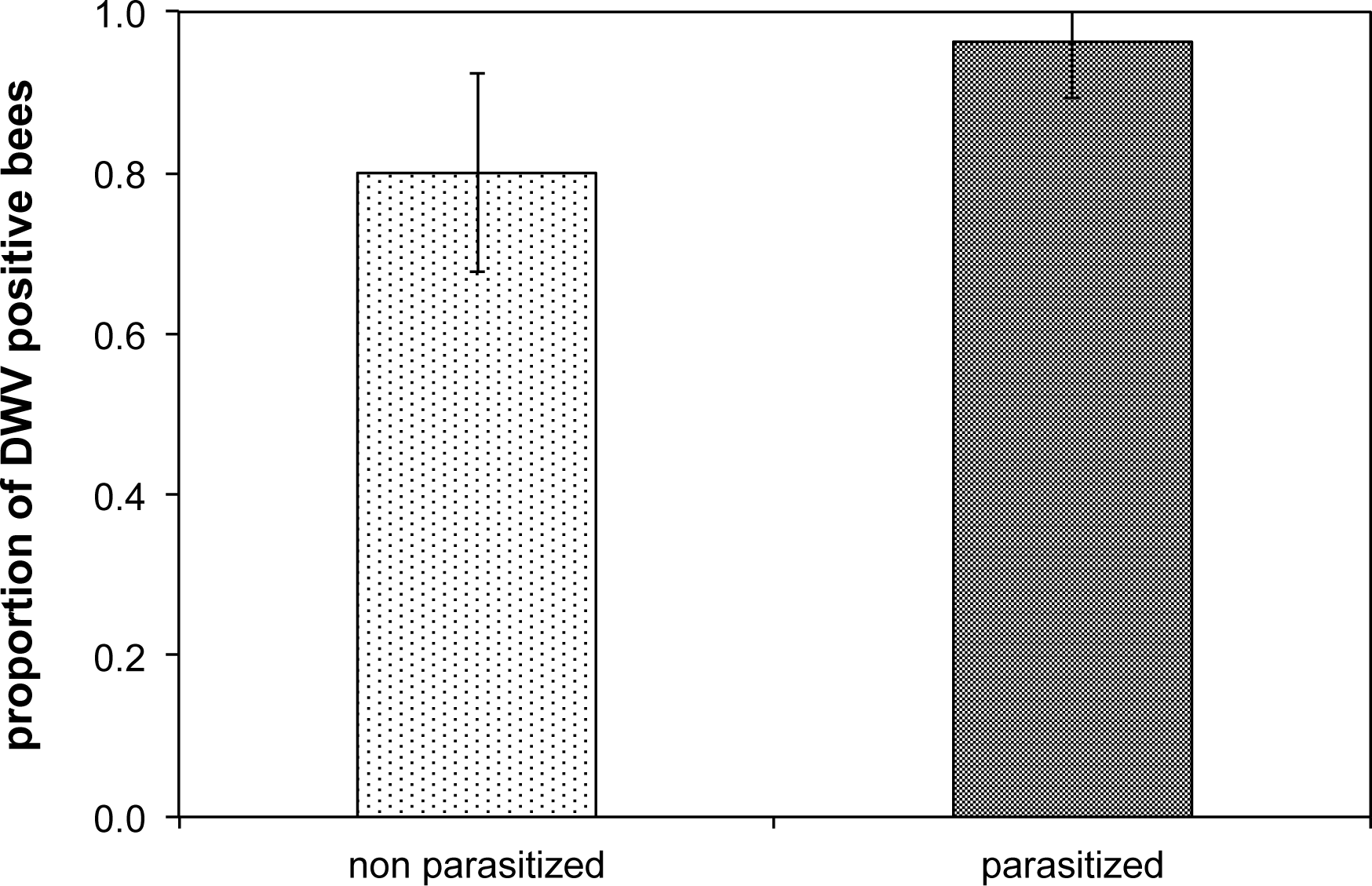
DWV prevalence in bees parasitized or not by a DWV infected mite. Error bars represent the estimated confidence limits of the proportion.

**Figure S3.**
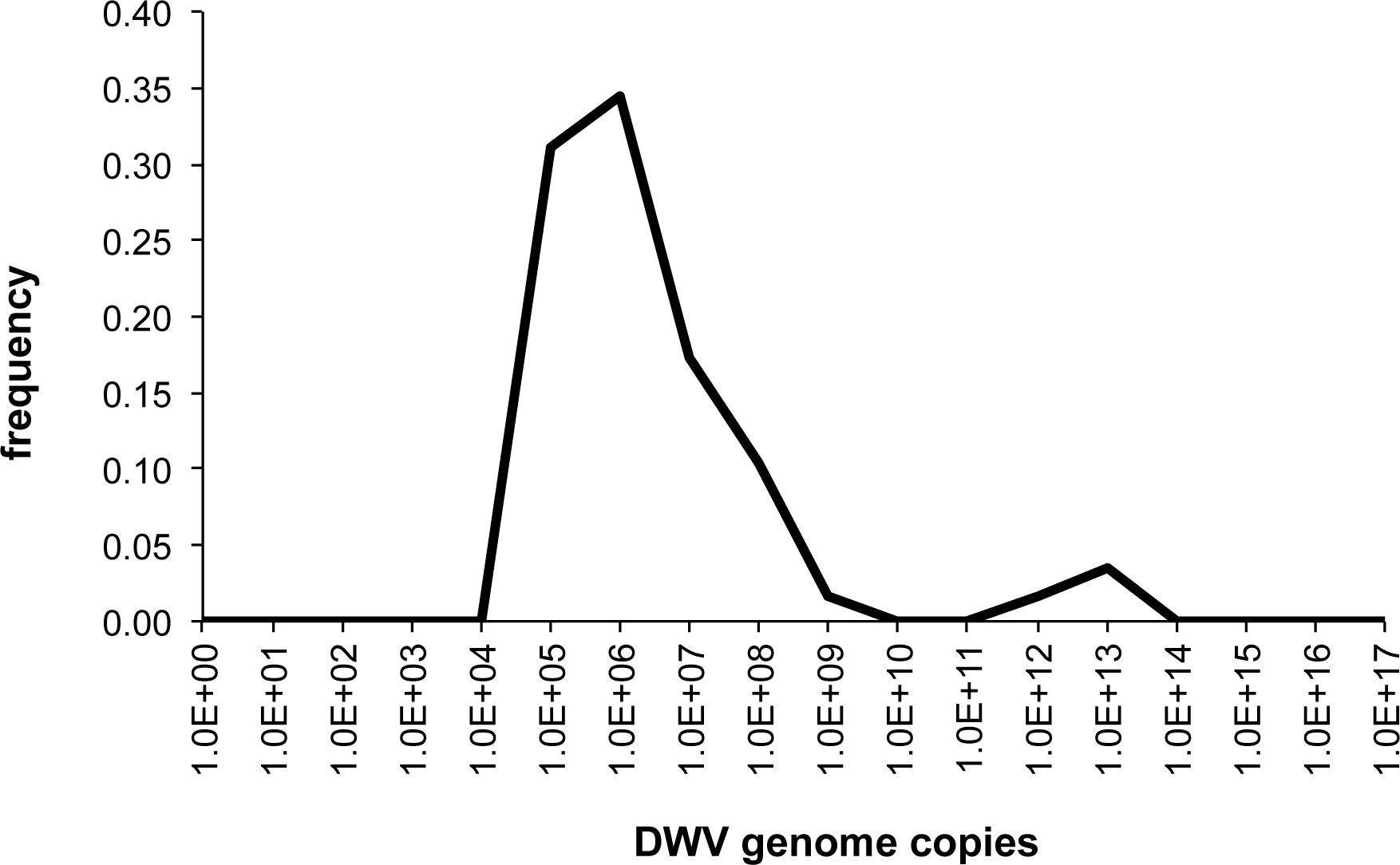
Frequency distribution of infection levels in bees parasitized with one mite in late summer.

**Figure S4.**
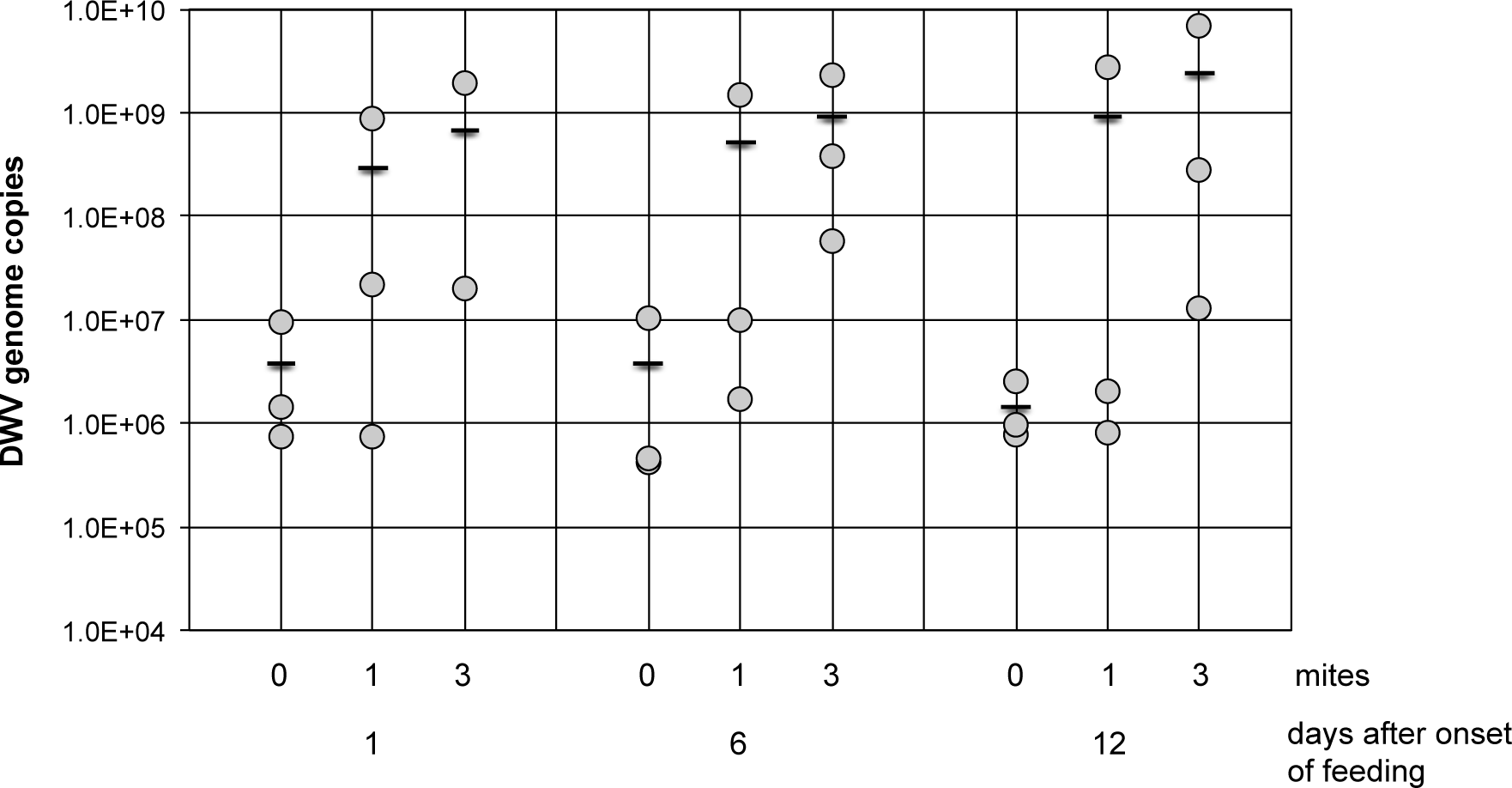
Viral load in individual bees artificially infested at the L5 stage with one, three or no mites and sampled after 1, 6 and 12 days (eclosion). A further sample, parasitized by 3 mites and collected 1 day after eclosion, with infection level=1.22E+03, does not appear on this graph.

**Figure S5.**
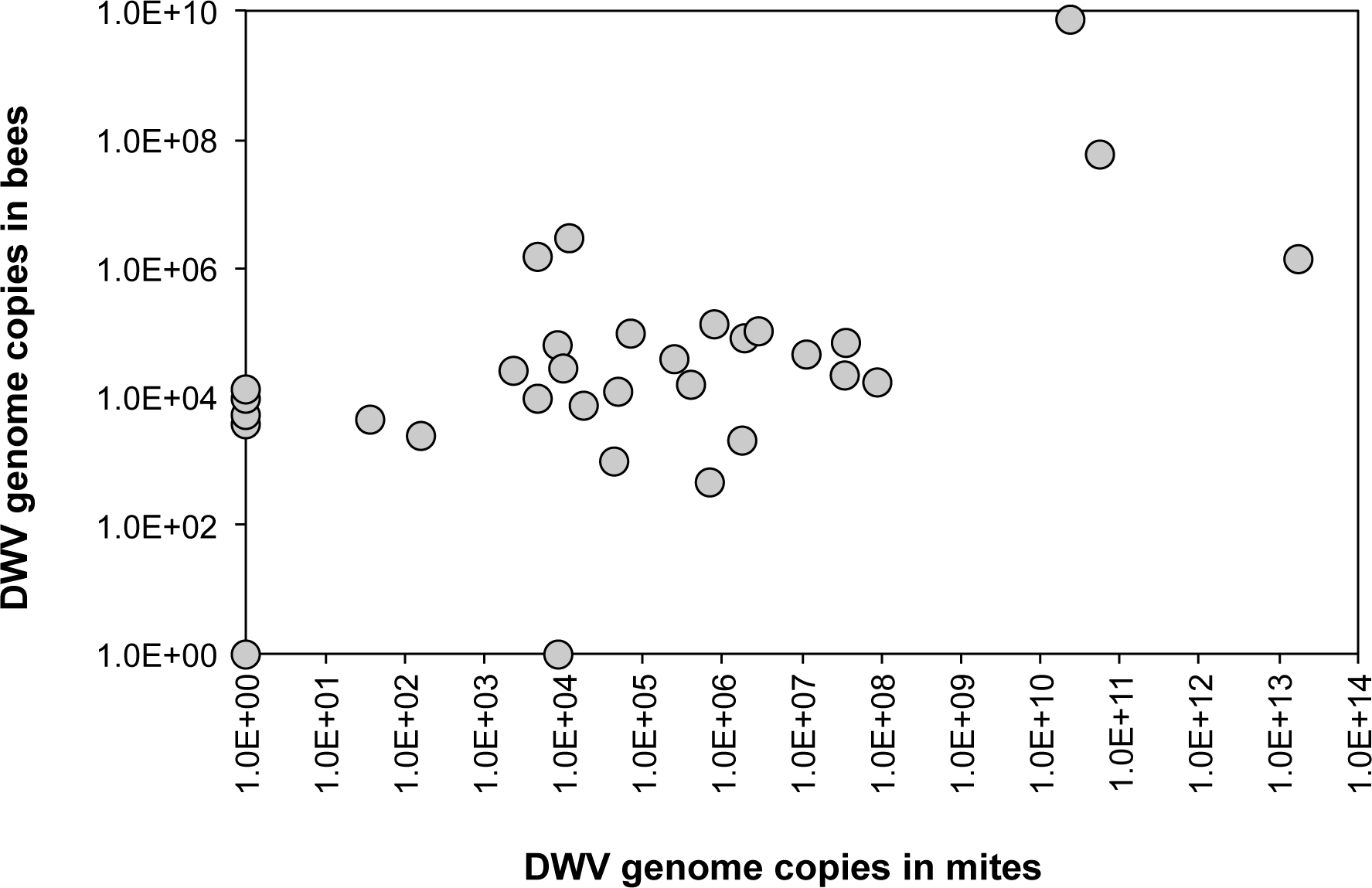
Infection level in infesting mites and infested bees. Note that zero values were transformed to 10.

**Figure S6.**
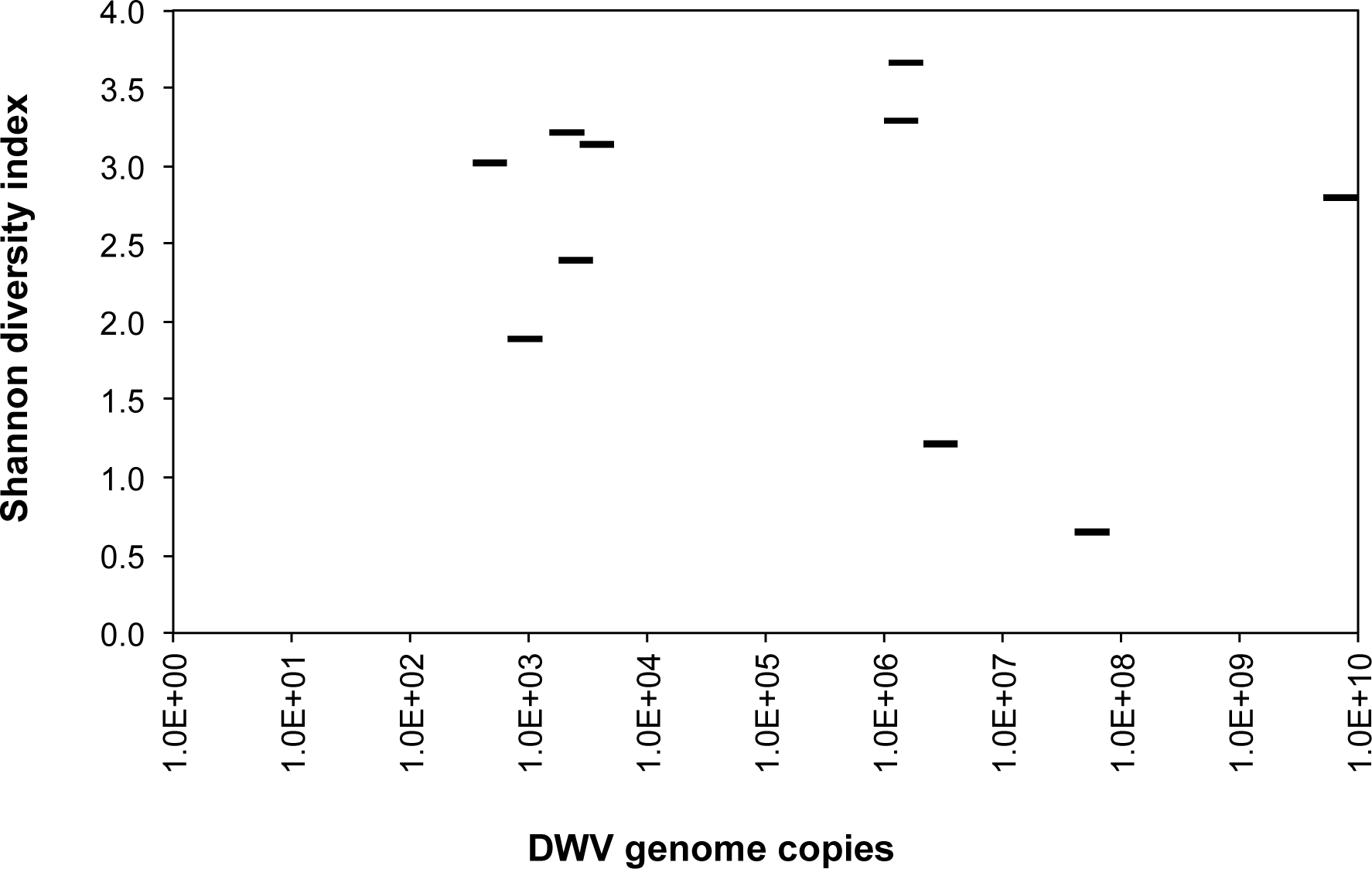
Diversity of the mutant cloud, as estimated with the Shannon index, and viral infection in bees.

**Figure S7.**
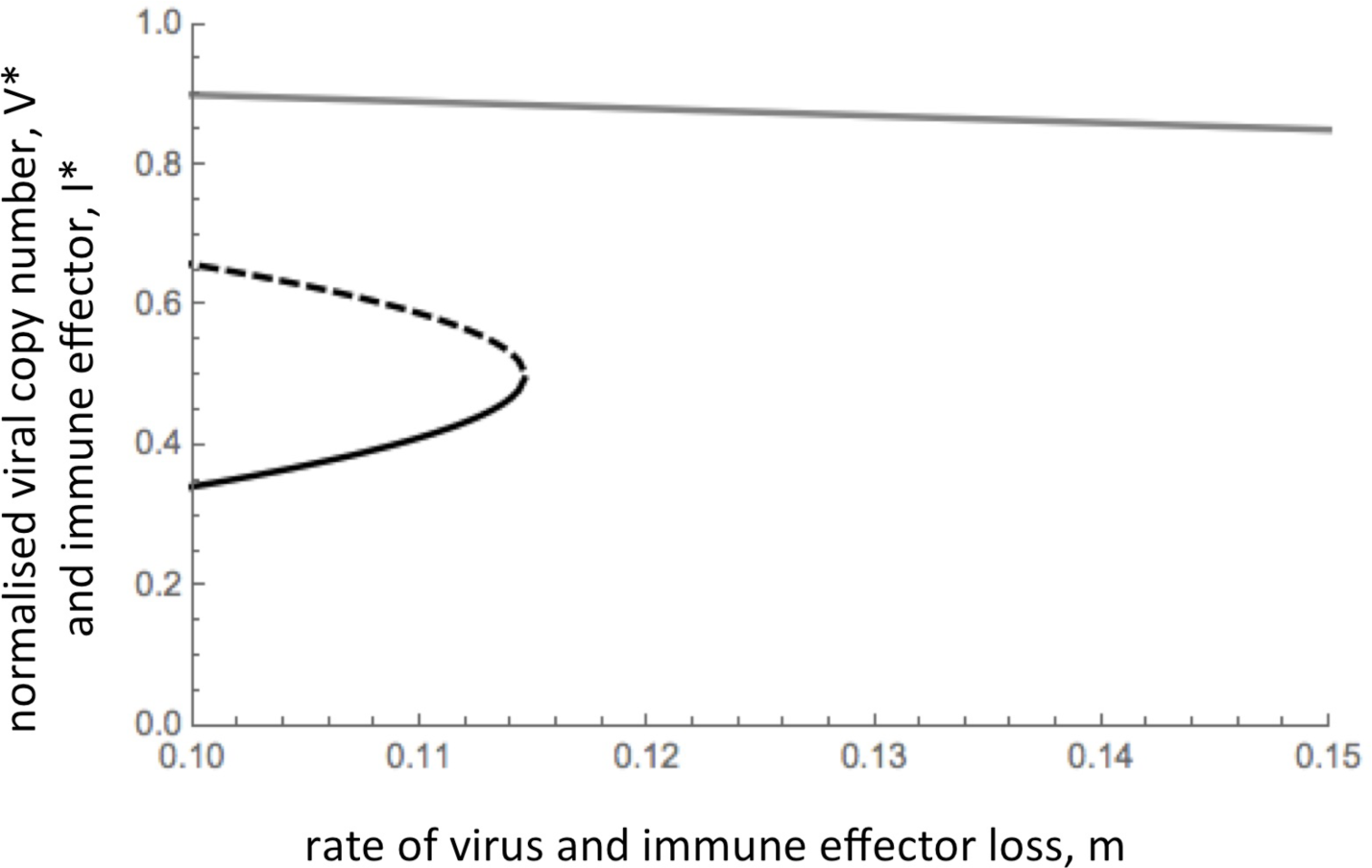
Increasing haemolymph loss destabilises viral dynamics. Stable (solid lines) and unstable (dashed line) equilibria are plotted for both viral titre (black lines, scaled to viral density that halts immune proliferation) and immune effector density (gray line, scaled to density that halts viral replication). Equilibrium expressions are defined in equations S4. Below the dotted line, the virus can be efficiently regulated by the immune-system to some intermediate (potentially cryptic) density, represented by the solid line. Above the dotted line (and for high *m*, any point to right of intersection with solid line), the virus cannot be efficiently regulated and a viral explosion ensues. A constant loss of viral particles and circulating immune effectors caused by mite feeding (increasing *m*, moving right along the solid lines) will first cause a gradual increase in copy number, V* (black line) and a correspondent decrease in circulating immune effectors, I* (gray line), and then at a defined point (intersection of solid and dotted lines), a viral explosion will ensue. Parameters are *x* = 0.09, *y* = 0.1 and *z* = 0.4.

**Figure S8.**
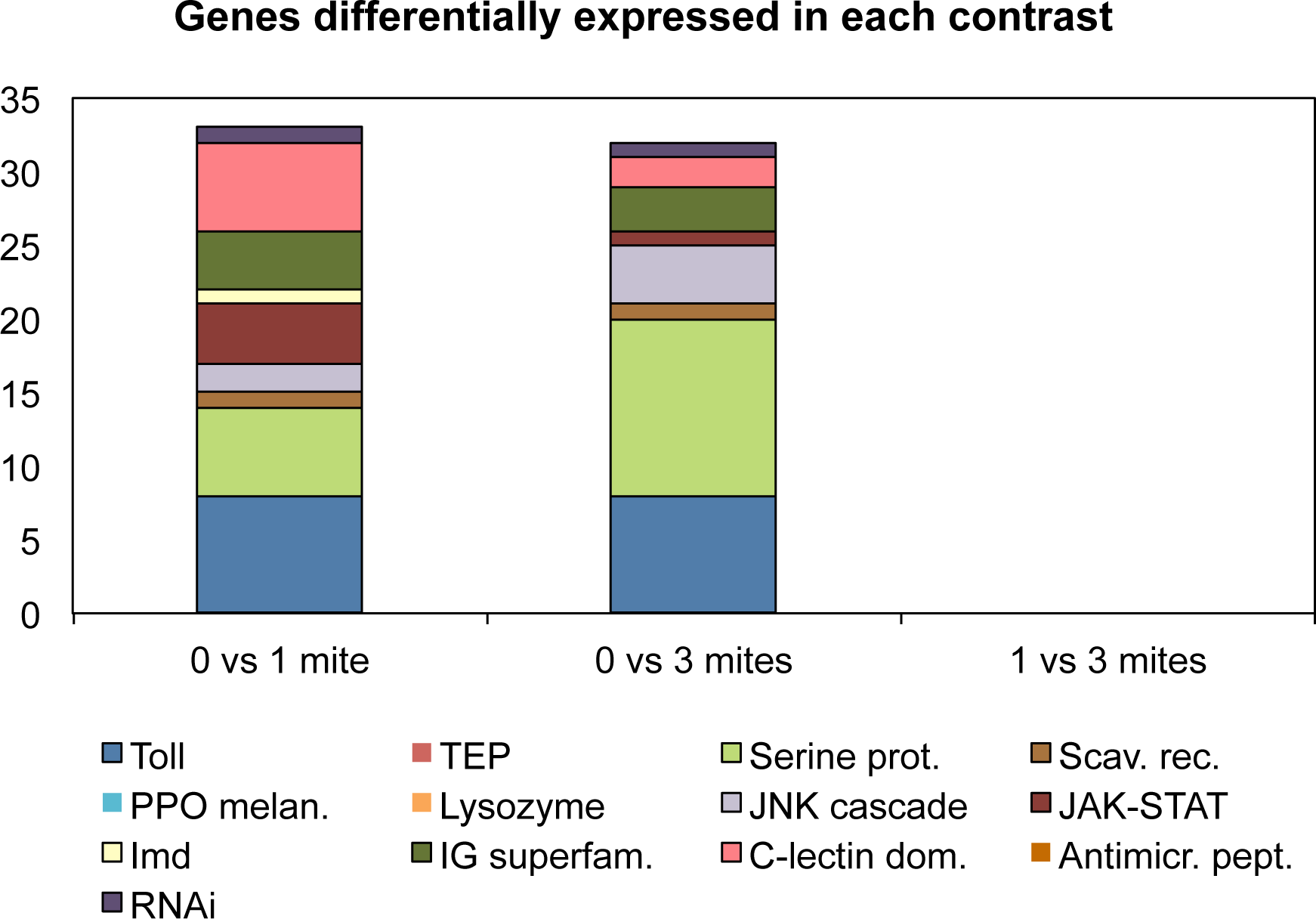
Differential immune response in bees infested by no, one or three mites. For each contrast between treatments the number of differentially expresses genes belonging to the canonical immune pathways is reported.

**Figure S9.**
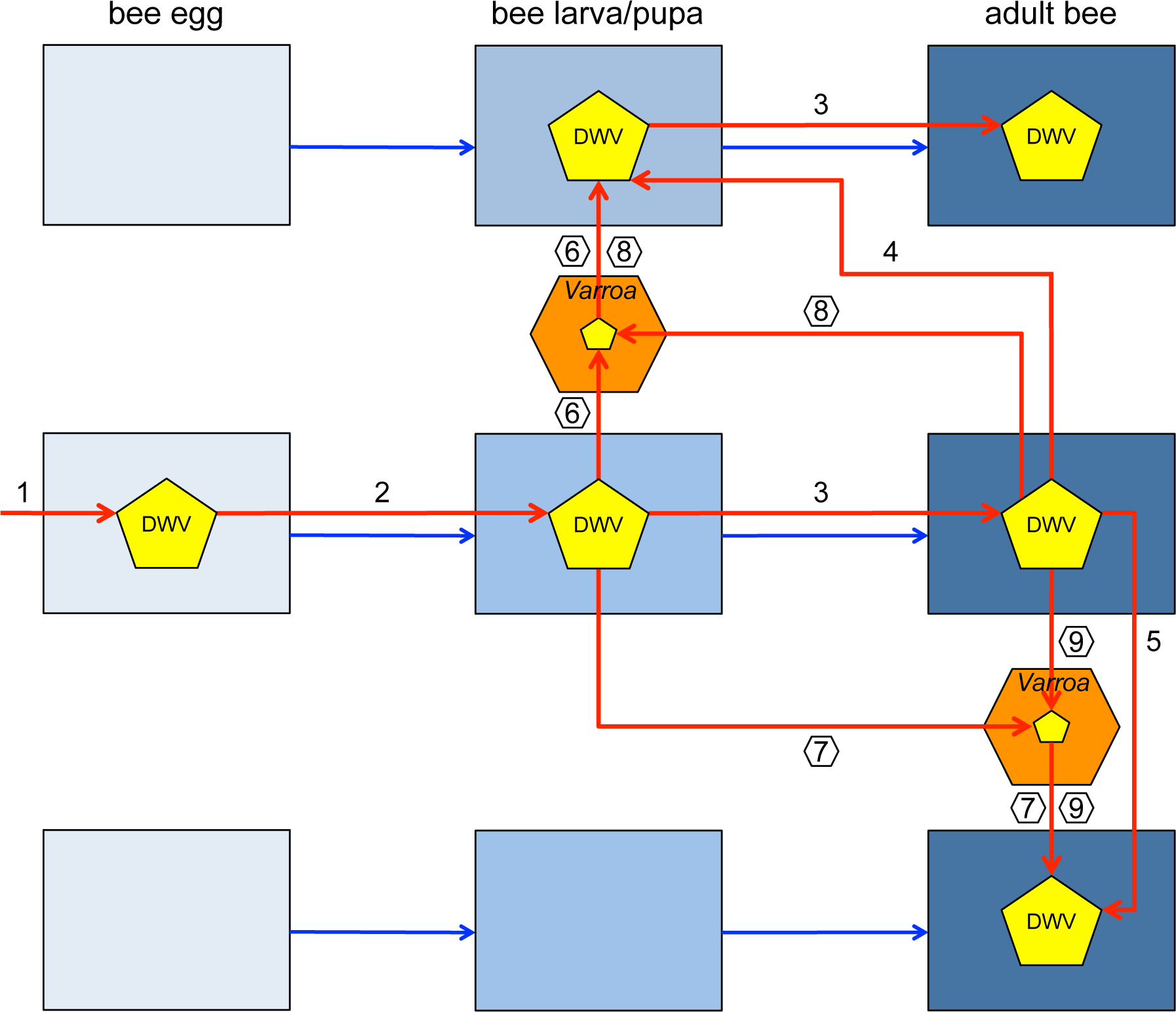
Possible routes of infection of DWV. The bees are represented by squares of different colours according to the developmental stage, the virus is a yellow pentagon and the mite an orange hexagon. Both trans-ovarial (from queen to egg, 1) and trans-stadial (from egg to larva, 2, and from larva to adult, 3) as well as horizontal transmission between nurse bees and larvae (4) and from adult to adult (5) are possible among bees. The *Varroa* mite can transfer the virus from pupa to pupa (6), pupa to adult (7) and viceversa (8) and adult to adult (9). Note the possible infection routes of the bee larva (2, 4), that are possible independently of the mite’s presence. The contribution of the mite as a virus vector is highlighted by transmission routes 6, 7, 8, 9.

**Figure S10.**
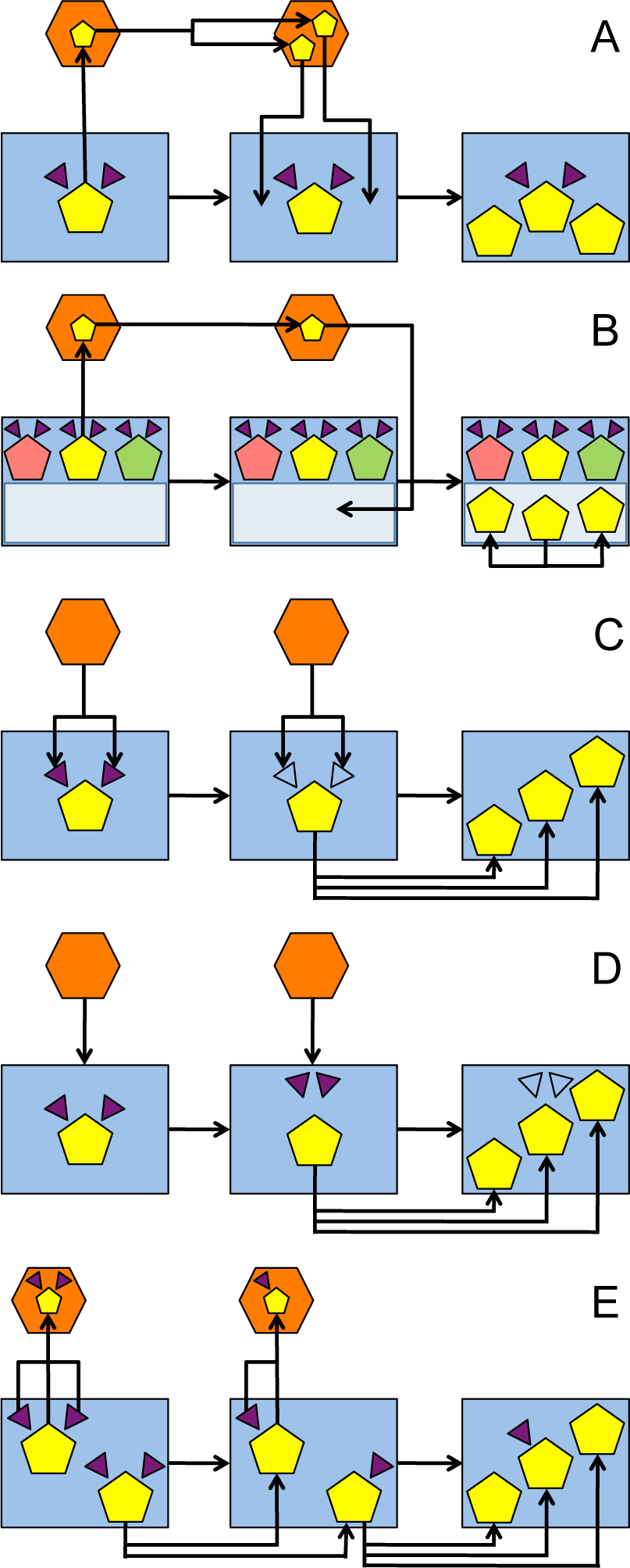
Current hypotheses about the role of the mite in facilitating viral infections. The bees are represented by squares, the virus is a yellow pentagon and the mite an orange hexagon; violet triangles (empty after suppression) represent the immune effectors. (A) The virus replicates within the mite and subsequently infects the pupa (8). (B) The mite favours the proliferation of a virulent strain of the virus (in the case represented here as an example, by injecting the pathogen into the haemolymph where replication is easier) (12, 13). (C) The mite suppresses the bee’s immune response (10). (D) The mite engages the same factors needed to sustain the antiviral response, releasing the pathogen from immune control (9). (E) The mite feeds upon the virus-contaminated haemolymph, subtracting both DWV and immune effectors, altering the dynamics of the systems, resulting in increased virus abundance (this article).

